## Supplementary Information for "Sequence-Based Generative AI-Guided Design of Versatile Tryptophan Synthases"

**The PDF file includes:**

Supplementary Figs. 1 to 7

**Other Supplementary Materials for this manuscript include the following:**

Supplementary Data 1.csv

Description: Sequences of all natural, evolved, and generated TrpBs used in this study, along with their sources.

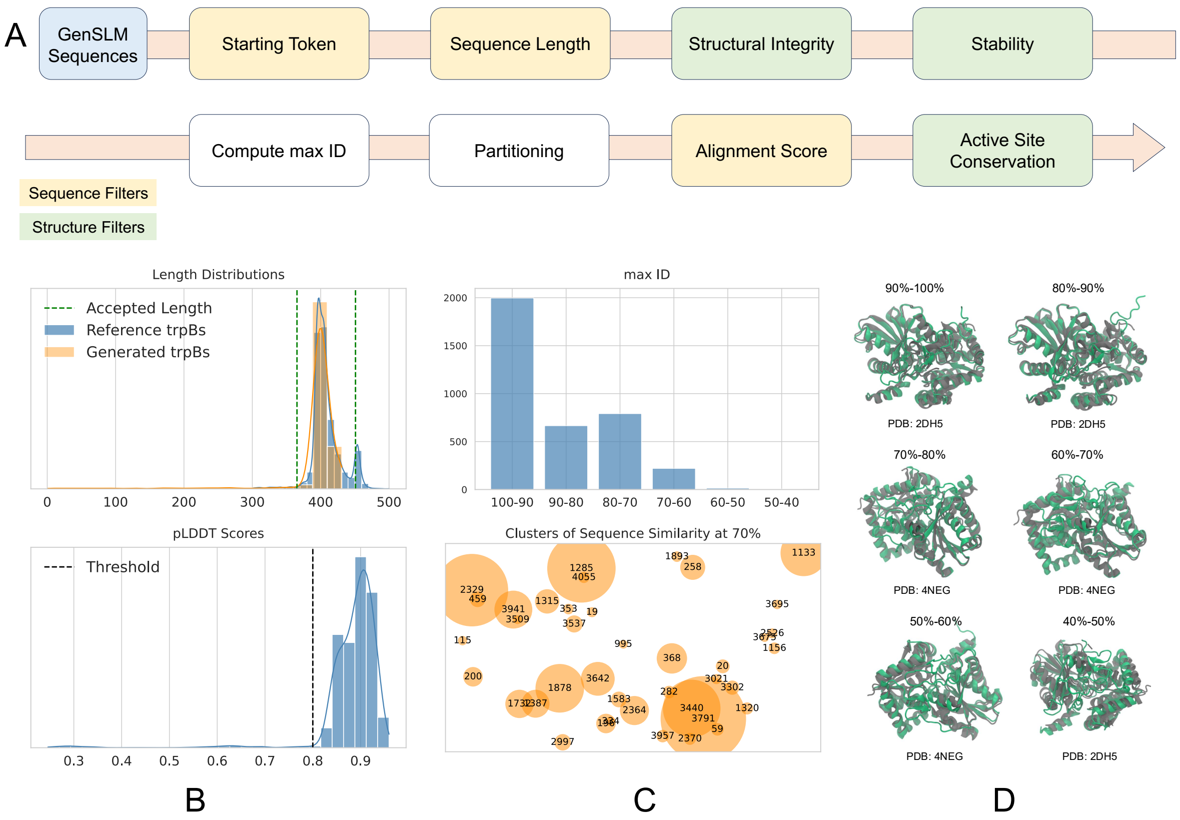


Figure S1: **Filtering of the generated *trpB* sequences**. (A) A schematic depiction of our proposed filtering process. (B) Distributions of length and pLDDT scores. (C) The sequence identity and sequence similarity of the generated *trpB* sequences. (D) The predicted structure of six selected samples from each partition of the max ID aligned with their closest match from the references.

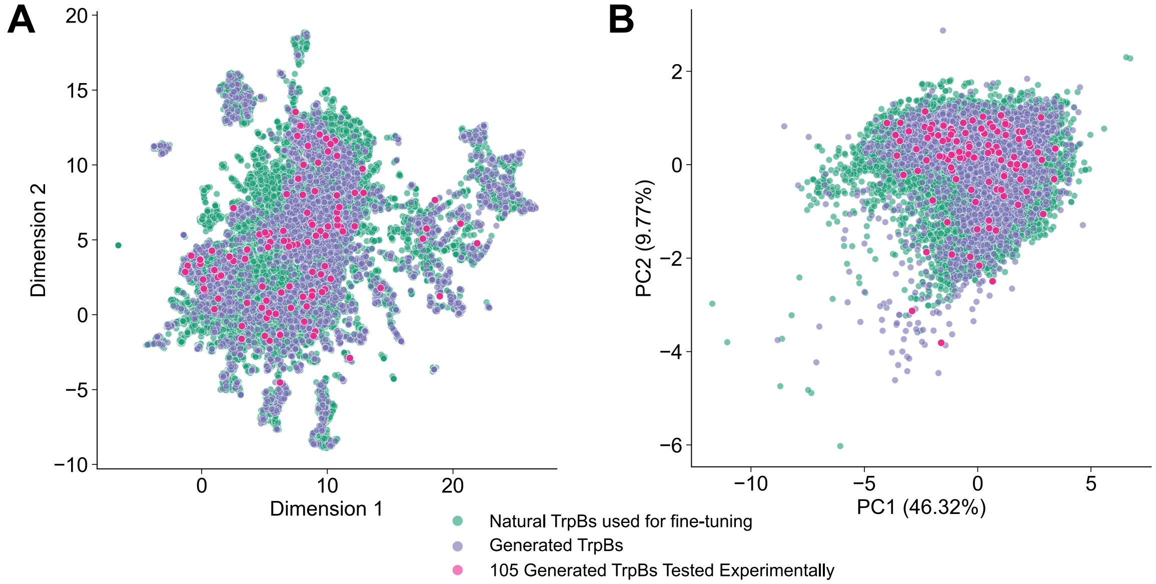


Figure S2: **2D visualizations of the GenSLM embeddings.** (A) UMAP projection of natural and GenSLM-TrpB sequences. (B) PCA projection of the same dataset.

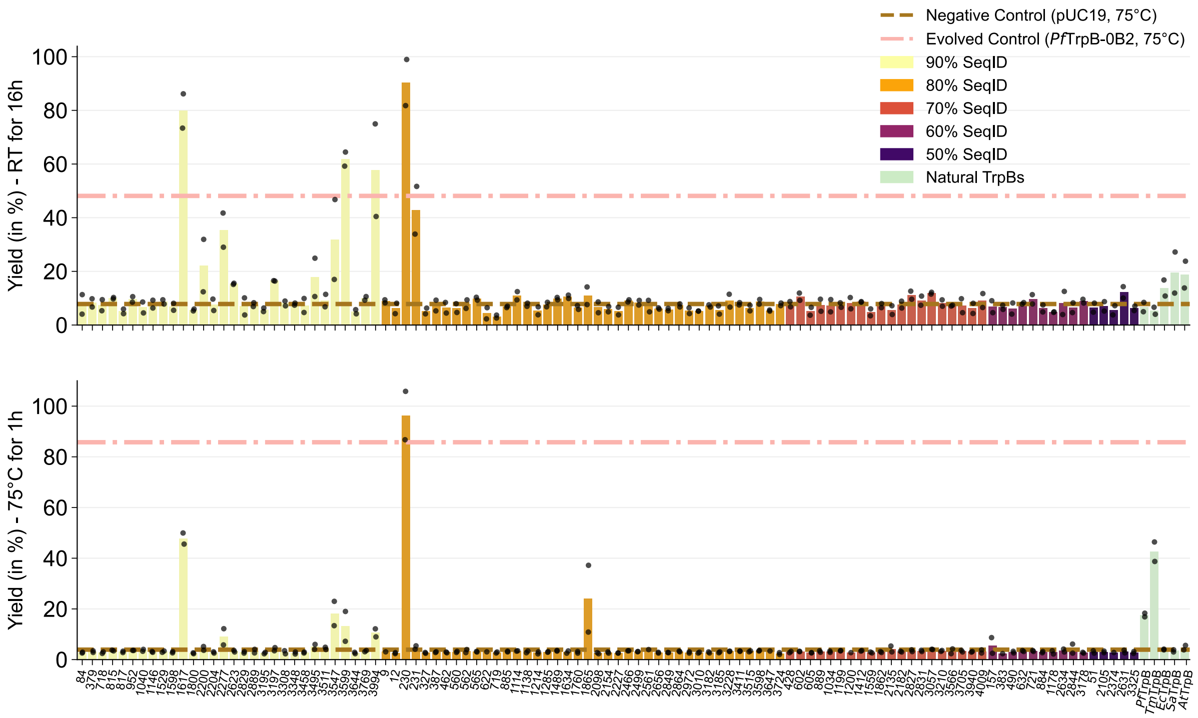


Figure S3: Tryptophan yields of GenSLM-TrpBs and natural TrpBs measured after 16 h at room temperature and 1 h at 75 °C. SeqID: sequence identity.

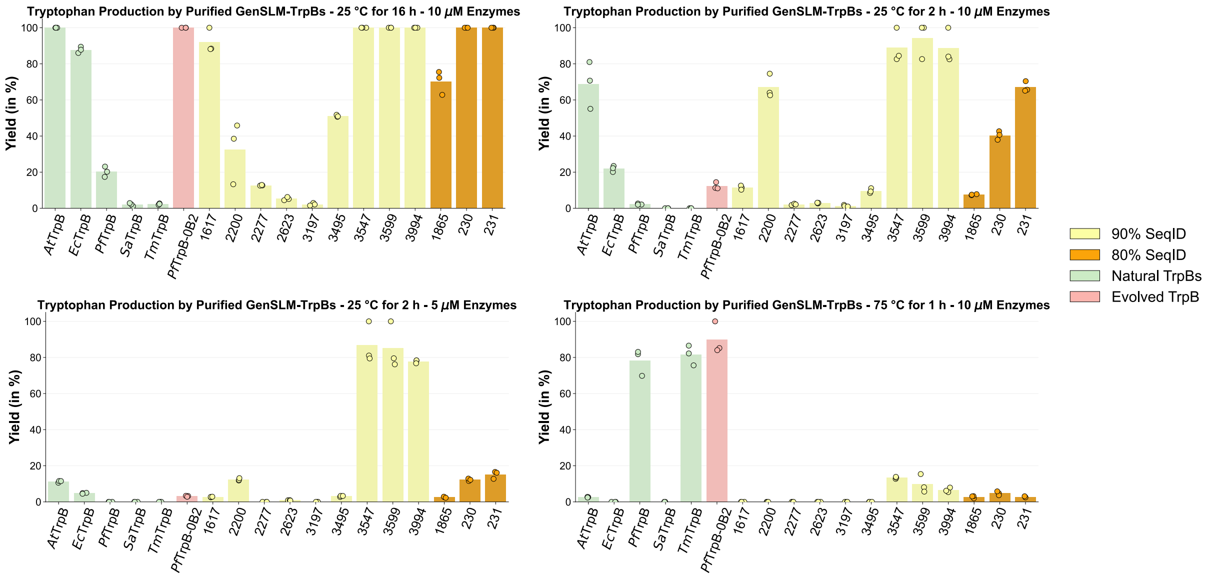


Figure S4: **Product yields of purified GenSLM-TrpBs.** Product yields of purified GenSLM-TrpBs measured under varying protein concentrations, reaction times, and temperatures in triplicates.

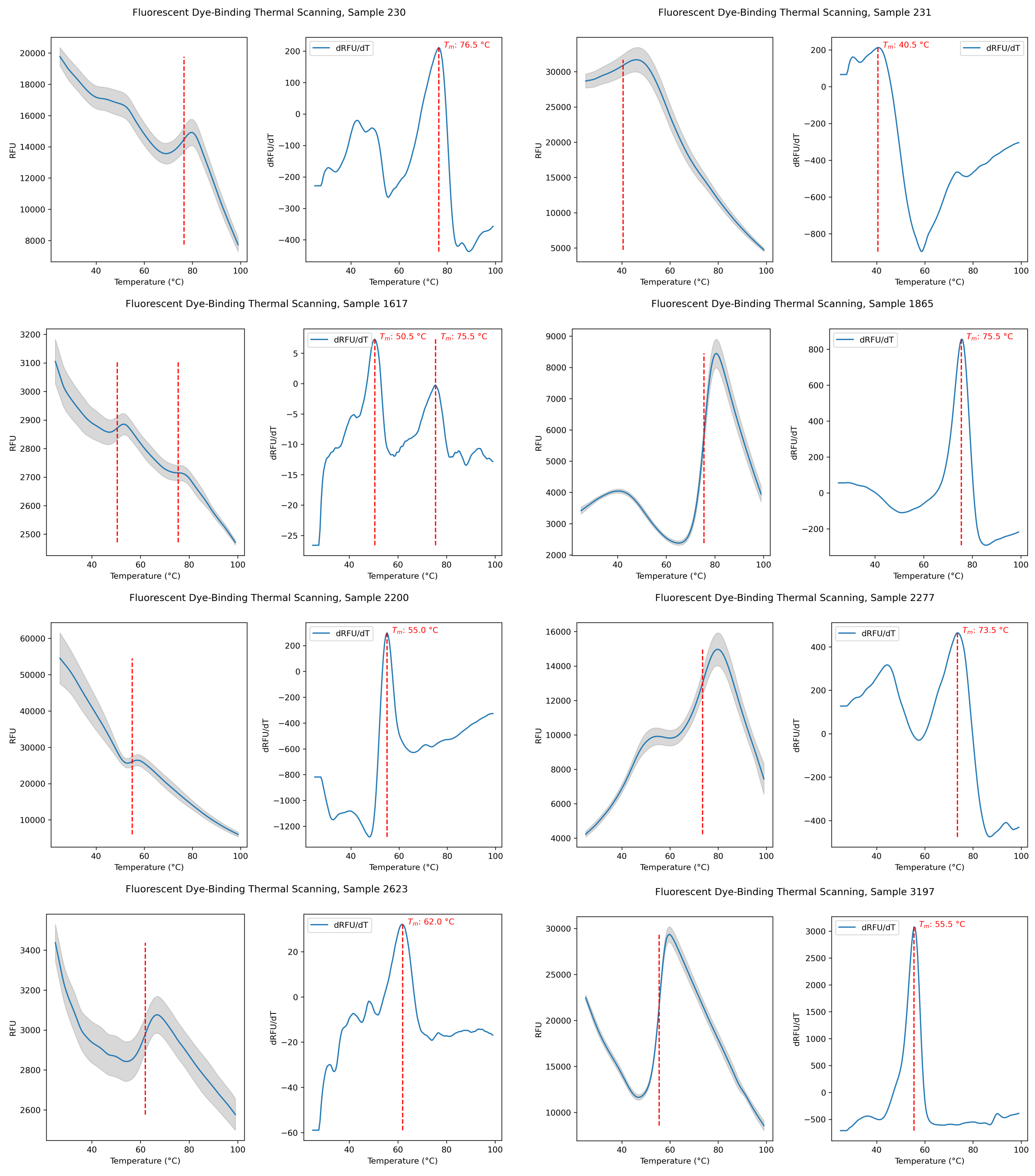


Figure S5: Melting temperature determined by thermoshift assay using purified GenSLM-TrpBs in triplicates.

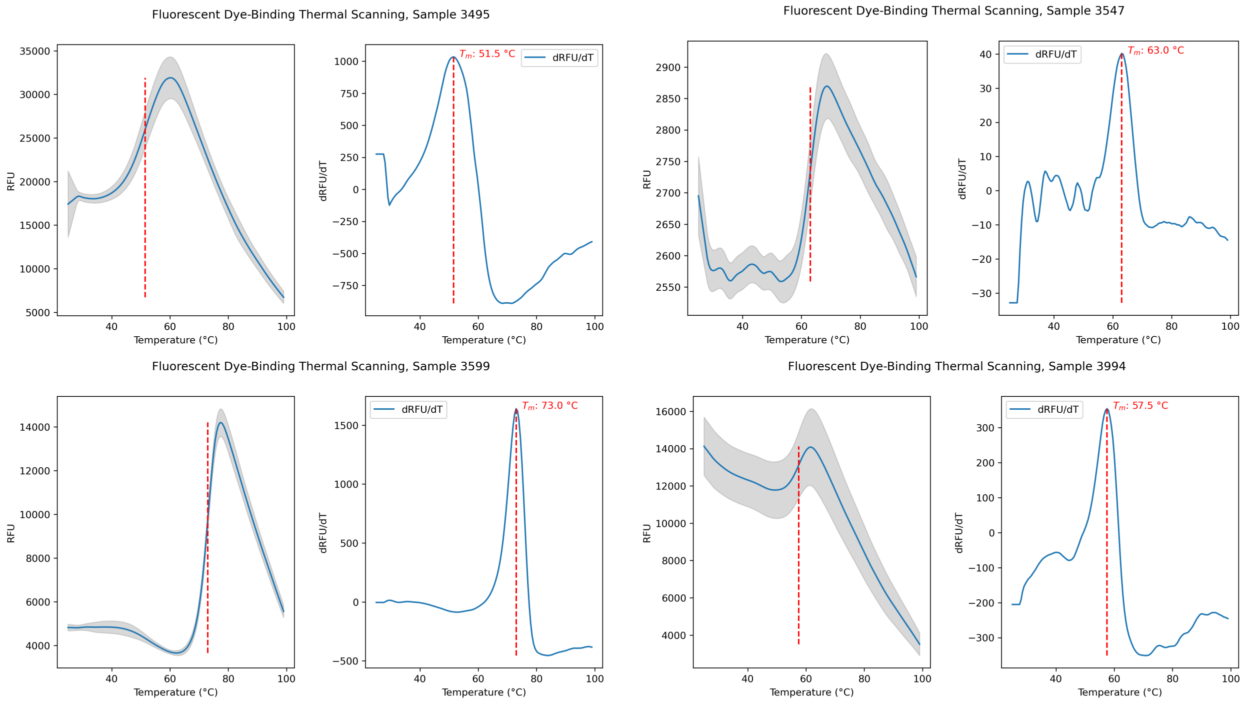


Figure S5: (continued): Melting temperature determined by thermoshift assay using purified GenSLM-TrpBs in triplicates.

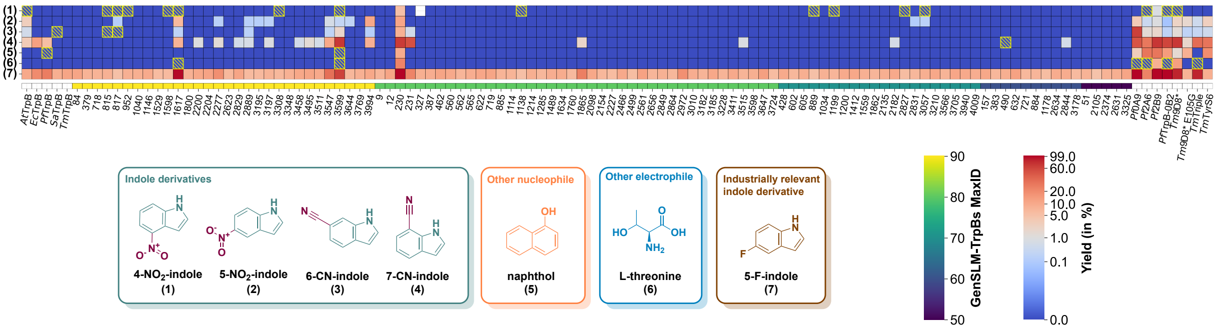


Figure S6: Yields of GenSLM-TrpBs with non-natural substrates estimated from absorbance at the isosbestic point (277 nm). Yellow dashed boxes denote reactions where product formation was confirmed by mass spectrometry above background levels, but remained below the UV detection threshold. Yields are displayed using a power-law normalization with $\gamma = 0.15$.

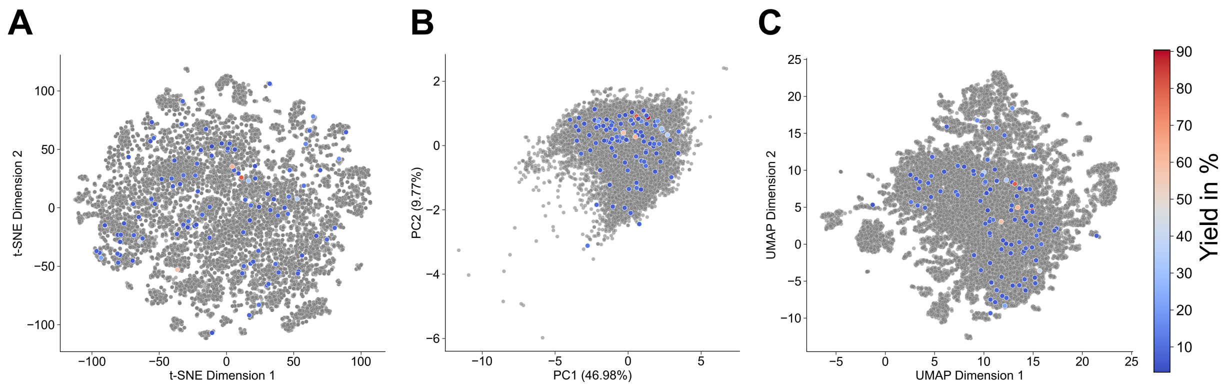


Figure S7: **2D visualizations of the GenSLM embeddings, colored by yield of tested variants.** (A) t-SNE projection. (B) PCA projection. (C) UMAP projection. Active GenSLM-TrpBs are distributed throughout the natural sequence space, indicating that the model did not converge to a single solution.
